# Extending causality tests with genetic instruments: an integration of Mendelian Randomization and the Classical Twin Design

**DOI:** 10.1101/134585

**Authors:** Camelia C. Minică, Conor V. Dolan, Dorret I. Boomsma, Eco de Geus, Michael C. Neale

**Author notes:** Corresponding author: Dr. Camelia C. Minicã, Vrije Universiteit Amsterdam, Department of Biological Psychology, Transitorium 2B03, Van der Boechorststraat 1, 1081 BT, Amsterdam, The Netherlands.

## Abstract

Mendelian Randomization (MR) is an important approach to modelling causality in non-experimental settings. MR uses genetic instruments to test causal relationships between exposures and outcomes of interest. Individual genetic variants have small effects, and so, when used as instruments, render MR liable to weak instrument bias. Polygenic scores have the advantage of larger effects, but may be characterized by direct pleiotropy, which violates a central assumption of MR.

We developed ***the MR-DoC twin model*** by integrating MR with the Direction of Causation twin model. This model allows us to test pleiotropy directly. We considered the issue of parameter identification, and given identification, we conducted extensive power calculations. MR-DoC allows one to test causal hypotheses and to obtain unbiased estimates of the causal effect given pleiotropic instruments (polygenic scores), while controlling for genetic and environmental influences common to the outcome and exposure. Furthermore, MR-DoC in twins has appreciably greater statistical power than a standard MR analysis applied to singletons, if the unshared environmental effects on the exposure and the outcome are uncorrelated. Generally, power increases with: 1) decreasing residual exposure-outcome correlation, and 2) decreasing heritability of the exposure variable.

MR-DoC allows one to employ strong instrumental variables (polygenic scores, possibly pleiotropic), guarding against weak instrument bias and increasing the power to detect causal effects. Our approach will enhance and extend MR’s range of applications, and increase the value of the large cohorts collected at twin registries as they correctly detect causation and estimate effect sizes even in the presence of pleiotropy.

## INTRODUCTION

Establishing causality in observational studies is important as knowledge of the relationship between a putative causal factor (exposure) and a potential outcome may inform rational treatment and prevention policies. While randomized controlled trials (RCTs) are the acid test of causality, they are expensive, time consuming, and may be practically or ethically unfeasible. For example, one cannot assign randomly individuals to a ‘low serum cholesterol condition’ in studying the causal effects of low serum cholesterol levels on cancer. An important alternative to the RCT is Mendelian Randomization (Katan ^1^, 2004).

Mendelian Randomization (MR) offers some traction in addressing causality by using genetic variants as instrumental variables to detect the causal effect of a modifiable risk factor (exposure) on a disease outcome in non-experimental settings ^2,3^. MR is quickly becoming the dominant approach to establishing causality; many recent applications have been published ^2,4-10^. The ascendency of MR is due to: dramatic drop in DNA genotyping costs, which has given rise to large genotyped samples, robust genetic associations established in genome-wide association studies (GWASs) ^11^, and the inherent advantages of MR, which include ecologic validity, robustness to reverse causation (from exposure to instrument) ^12^ and confounding ^13^.

MR requires instruments with a relatively strong direct relationship with the exposure. A disadvantage of many genetic variants is that they have weak effects ^14^. Weak instruments confer insufficient statistical power, and render MR liable to weak instrument bias ^13,15,16^. Combining the weak genetic effects into a polygenic risk score (PGS) is a possible route to increase the strength of the genetic instrument ^13,17-19^. However, the MR assumption that the instrument is not pleiotropic (has no direct effect on the outcome) is stronger in the case of a PGS instrument ^13,16,20,21^. A PGS comprises many genetic variants, any of which may directly affect both the exposure and the outcome, or may include variants in linkage disequilibrium with variants affecting the outcome. As demonstrated by twin studies ^22-27^ and by polygenic risk score analyses ^28-30^, many genetic variants associate with multiple phenotypes, suggesting pervasive pleiotropy ^31-35^.

Several methods are currently in use as means to tackle the ‘no pleiotropy’ assumption. Some approaches apply prior selection criteria to increase the probability that the instruments are valid. For instance, the stepwise procedure implemented in the R-package gtx ^36^ employs iteratively a heterogeneity test to discard from a polygenic score genetic instruments yielding significant heterogeneity in the estimates of the causal effect. Possible heterogeneity is assumed to be indicative of pleiotropy. The efficiency of this method depends on its power ^36^ to detect heterogeneity arising from pleiotropy. However, heterogeneity may arise from sources other than pleiotropy, so that one may needlessly weaken the instrument by removing valid genetic variants. Johnson ^36^ noted that the performance of this procedure in terms of bias and type I error rates within the MR context has yet to be established. Other approaches like those based on the median estimator ^37^ can handle up to 50% invalid instruments. However, the strong MR assumption still applies to the variant(s) yielding the median causal effect. MR-Egger regression ^38^ is another approach that, with weaker assumptions, gives consistent estimates even when all instruments are pleiotropic. The estimator is (asymptotically) consistent under the weaker assumption that the effect of the instrument on the exposure is uncorrelated with the effect of the instrument on the outcome (i.e., the Instrument Strength Independent of Direct Effect assumption). However, the InSIDE assumption is unlikely to hold universally ^38^, as shared molecular mechanisms (i.e., direct/biological pleiotropy) are expected to yield correlated effects on the associated traits (see, for examples, ^34^). Furthermore, Bowder et al. noted ^38^ that there are other plausible paths from the instrument to the outcome (except direct paths, or indirect, via the exposure), for example, via confounders affecting both traits, or due to linkage disequilibrium between the instrument and a genetic variant affecting the outcome. In this case the estimate of the estimates of the causal effect in Egger regression will be biased ^39^. Finally, although MR-Egger uses multiple genetic variants to estimate the causal effect, these instruments are employed individually (i.e., not combined in a polygenic score), and so the approach (like the median-based approaches) remains liable to weak instrument bias ^40^.

The utility of the classical twin design (CTD) in the study of direction of causality is well established ^41-45^. The present aim is to combine MR and CTD into a single model. A similar approach was proposed by Kohler et al. who integrated CTD with the instrumental variable method ^44^. Our focus is on combining MR with CTD to render testable the ‘no pleiotropy’ assumption. Particularly, we address issues of identification and statistical power associated specifically with a (poly-)genic instrument, which may be related directly to the outcome (i.e., violating the no-pleiotropy assumption). Integrating MR with CTD has three advantages: 1) it allows one to relax the strong MR assumption concerning the instrument’s conditional independence of the outcome (conditional on the exposure and confounders, i.e., the no pleiotropy assumption; 2) by accounting for pleiotropy, the approach facilitates the use of PGS as instruments; and 3) in specific circumstances, the approach confers substantial gains in power relative to the standard MR approaches.

## METHODS

Direction of Causation (DoC) twin model was advanced as an exploratory approach to establish direction of causation between two correlated traits ^43,46-49^. In contrast, MR ^3,13^ is used to test unidirectional causation (from designated causal exposure to outcome). Here we propose the MR-DoC twin model, developed by imposing restrictions on the DoC parameters to represent unidirectional causality hypotheses, and by extending the model to include measured genetic variants as instrumental variables. Integrating MR with DoC allows us to test certain MR assumptions.

### The Direction of Causation twin model

The Direction of causation (DoC) twin model ^43,46-49^ uses cross-sectional data observed in monozygotic (MZ) and DZ twins to test causal hypotheses regarding the association between two traits. In contrast to MR, DoC does not necessarily involve a prior hypothesis concerning the causal direction. The path diagram of such a model is shown in Figure 1, given an exposure variable X and an outcome variable Y, observed in DZ twins.

**Figure 1:**
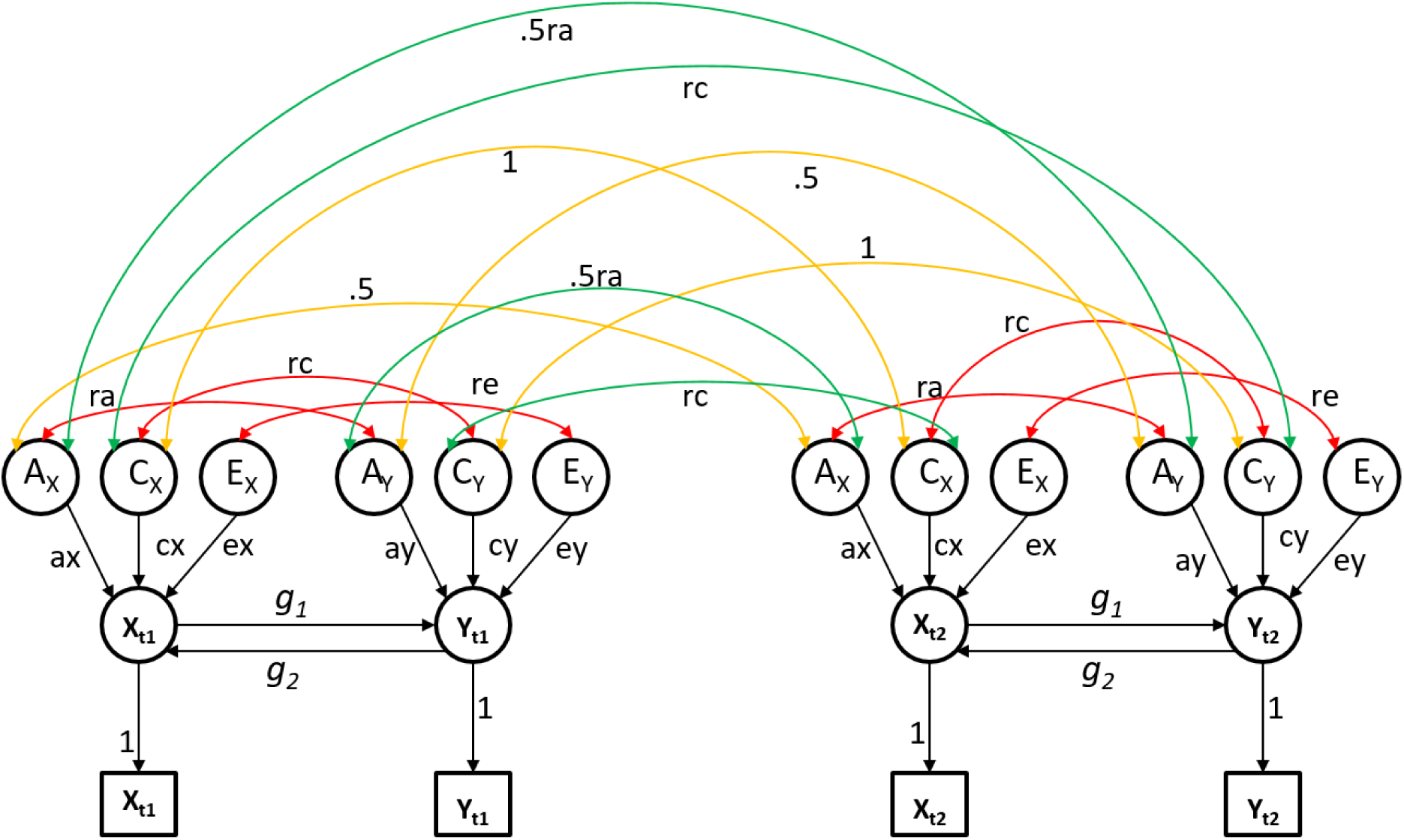
Path diagram representing the direction of causation twin model given two traits: variable X and variable Y measured in dizygotic (DZ) twins (t_1_ and t_2_). Squares represent observed variables, while circles represent latent variables. A, C and E stand for additive genetic effects, shared and unique environmental effects, respectively. The double headed arrows represent within/between twins covariances of additive genetic effects (ra), shared environmental effects (rc) and unique environmental effects (re). The cross-twin covariance between additive genetic effects is fixed to .5 (1) for DZ (MZ) twins. DZ (MZ) twins are expected to share on average 50% (100%) of the genetic effects underlying both traits, hence the cross-twin cross-trait covariance is fixed to .5(1)ra for DZ (MZ) twins.. Single headed arrows represent causal effects. Note, the model as depicted, is not identified. Typically ra, rc, and re are assumed to be zero in the application of the DoC twin model.

In Figure 1, X and Y are mutually causally related (parameters g_1_ and g_2_). The traits are subjected to the influence of latent additive genetic (A_X_ and A_Y_), shared (C_X_ and C_Y_) and unique (E_X_ and E_Y_) environmental effects, influences which can be direct or indirect, i.e., via the causal paths. As an instance of CTD, this model has the usual assumptions concerning random mating and the absence of genotype-environment interplay (GxE interaction, GxE covariance). The cross-twin correlation of the shared environmental variables is assumed to equal 1 within-trait, and *rc* across traits, regardless of zygosity. By definition, the cross-twin correlation of unique environmental effects is fixed to zero both within and across traits.

The model as depicted in Figure 1 is not identified; it requires additional restrictions to identify the parameters. By imposing restrictions on the parameters, one can model several alternative hypotheses concerning the observational association between X and Y. The ‘tertium quid’ hypothesis, that a third variable causes both traits, can be tested by constraining the parameters g_1_ and g_2_ to equal zero (i.e., the saturated bivariate model). Uni- and bidirectional causal hypotheses can be tested by fixing to zero the within- and cross-twin cross-trait genetic and environmental correlations (ra, rc, re), and estimating the causal parameters g_1_ and/or g_2_ (i.e., the uni- and the bidirectional causality models). These competing and nested alternative hypotheses can be tested by likelihood ratio ^42,43^, provided: 1) the two traits differ in their sources of variance ^43^; and 2) there are at least three sources of variance influencing the traits (i.e., A, E, and either C or D (dominance)) ^50^. Given the assumptions mentioned above, we have ^43,49^:

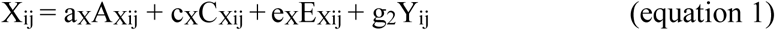

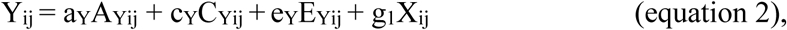

where subscript i stands for twin pair, and j for twin (j=1,2).

### Standard Mendelian Randomization

The MR model is an instrumental variable regression model, which employs genetic variants as instrumental variables to test causal hypotheses regarding the association between an exposure and an outcome ^3,13^. Here we assume that the instrument is a PGS. Three assumptions must hold for a genetic variant to be a valid instrumental variable, as shown in Figure 2. *Assumption 1*: The genetic instrument (PGS) is robustly associated with the exposure variable X (b_1_≠0 in Figure 2); *Assumption 2*: PGS is independent of confounders C (m=0; PGS ⊥ C); *Assumption 3*: PGS is independent of the outcome variable Y conditional on the exposure X and confounders C (b_2_=0; PGS ⊥ Y | X, C).

**Figure 2:**
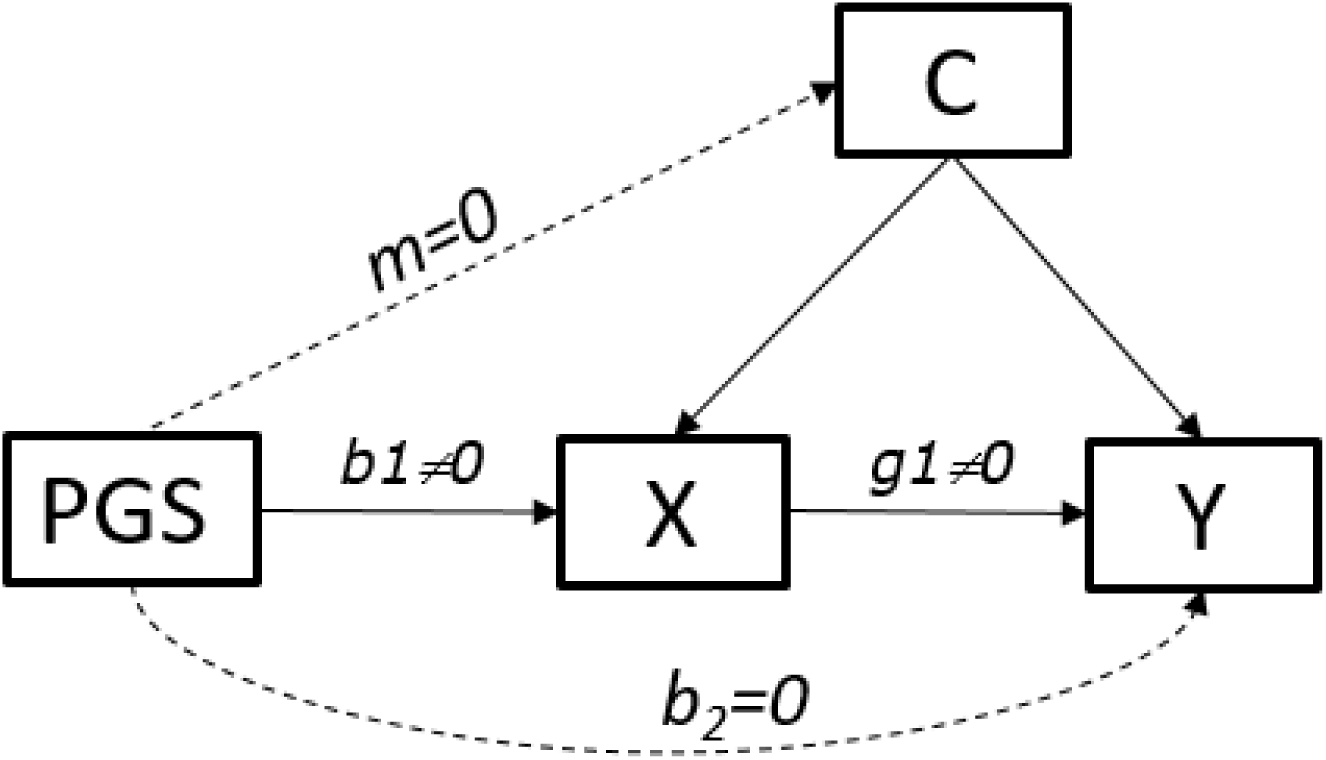
Assumptions in Mendelian Randomization. Hypothesis: X causes Y. By assumption, the regression coefficients (m and b_2_) associated with the dashed lines are zero. Abbreviations: PGS – polygenic score; X – exposure variable; Y – outcome variable; C – confounders; Abbreviations: PGS – polygenic score; X – exposure variable; Y – outcome variable; C – confounders;

In MR, the third assumption pertains to possible pleiotropic effects of the instrument (PGS), or to the likelihood of including variant(s) in linkage disequilibrium with variants affecting the outcome. In practice, the ‘no pleiotropy’ assumption may be violated, particularly when the instrument is a PGS combining the effects of multiple genetic variants (note that a single variant with pleiotropic effects in principle renders the polygenic score invalid as an instrumental variable). This core MR assumption is illustrated in Figure 3 where we consider several MR models with and without pleiotropic instruments, and pinpoint the definition of the no pleiotropy assumption.

**Figure 3:**
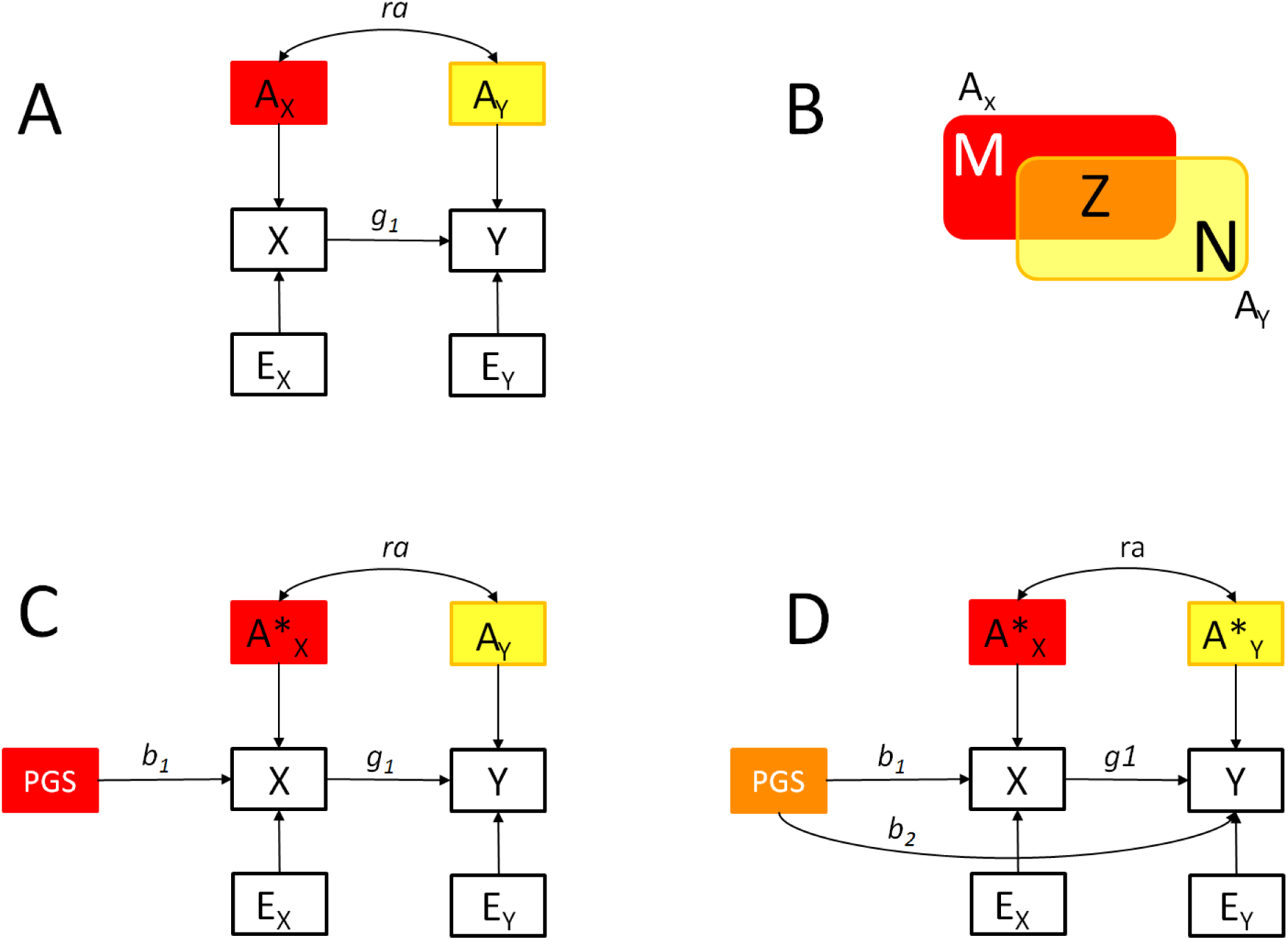
A) X directly influences Y (parameter g_1_). In addition, the additive genetic variables A_X_ and A_Y_ are correlated (parameter ra). B) The set of loci M underlies the variance of A_X_, but does not underlie the variance of A_Y_, i.e., in set theory notation, M=A_X_ /A_Y_. Likewise, N=A_Y_/A_X_, i.e., the set of loci N contributing to the variance of A_Y_ but not to the variance of A_X_. Z represents the intersection of A_X_ and A_Y_, that is, the set of loci Z underlies both A_X_ and A_Y_, i.e., Z=A_Y ⋂_ A_X_. Note that the set Z may contain pleiotropic loci, where the pleiotropy is due to direct effects or due to linkage disequilibrium; C) The MR model with a polygenic instrument (PGS) and ‘no pleiotropy’. PGS is associated with X (parameter b1), but is assumed to have no direct influence on the outcome Y. This model holds only if the instrument PGS is constructed on the basis of a subset of variants from set M. In the presence of PGS, A*x is a residual (in the regression of X on PGS). D) MR with pleiotropic genetic instrument. In this model, the PGS is constructed on the basis of a sample of genetic variants taken from set Z. The parameter b_2_ accommodates the fact that the set of variants used to construct PGS underlies the variance of both A_X_ and A_Y_. The no pleiotropy assumption implies b_2_=0.

Among the methods of causal effect estimation in the standard MR are the two-stage least squares (2SLS) and the ratio of coefficients. In 2SLS, first, the instrumental variable (e.g., the polygenic score) is used to predict the exposure X, and second, the outcome is regressed on the predicted values of X. In the ratio of coefficients method, the causal effect is computed as a ratio of regression coefficients, with the numerator obtained in the regression of the outcome on the instrument, and the denominator obtained in the regression of the exposure on the instrument ^13^. Both methods are based on least squares estimation, and are expected to yield equivalent results in MR studies involving a single instrumental variable ^13^.

The standard MR model can also be fitted in a single step as a structural equation model (as depicted in Figure 3 (panel C), with maximum likelihood (ML) estimation. The causal parameter 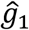 in Figure 3C can be tested by the means of a likelihood ratio or a Wald test.

### Integrating Mendelian Randomization with the Direction of Causation twin model (the MR-DoC model)

In observational studies, MR and DoC twin model offer some traction in testing a hypothesized direction of causality. As demonstrated below, the combined MR-DoC model has definite advantages over the individual approaches, in terms of power and assumptions. Figure 4 displays a path diagram of the MR-DoC model. Note that, the model as depicted is not identified. We consider the issue of identification below.

**Figure 4:**
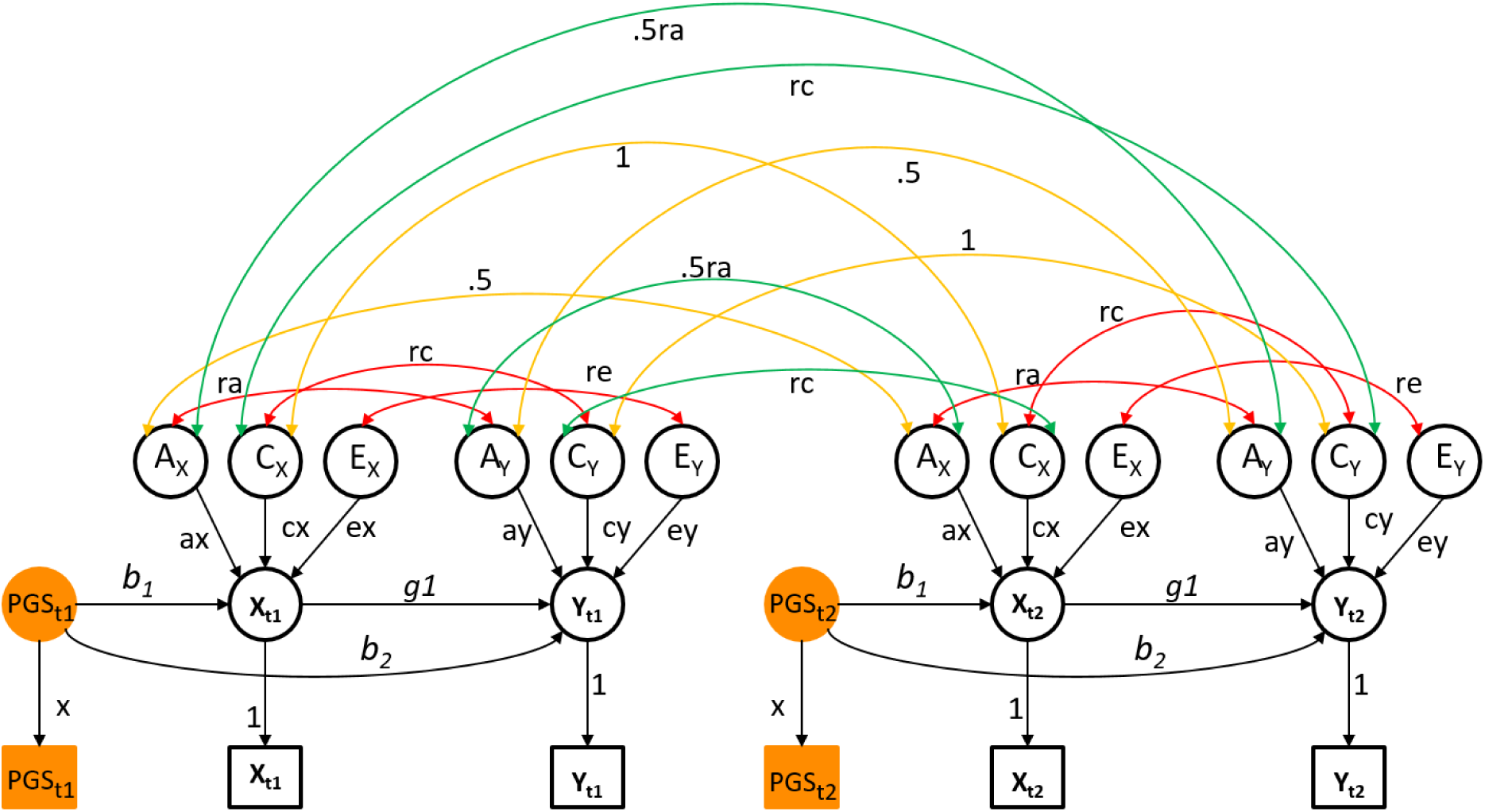
Path diagrammatic representation of the MR-DoC model in DZ twins. The parameter x equals the standard deviation of the observed instrument, i.e., PGS in the circle is standardized. The model as depicted is not identified (see Table 1).

**Table 1:**
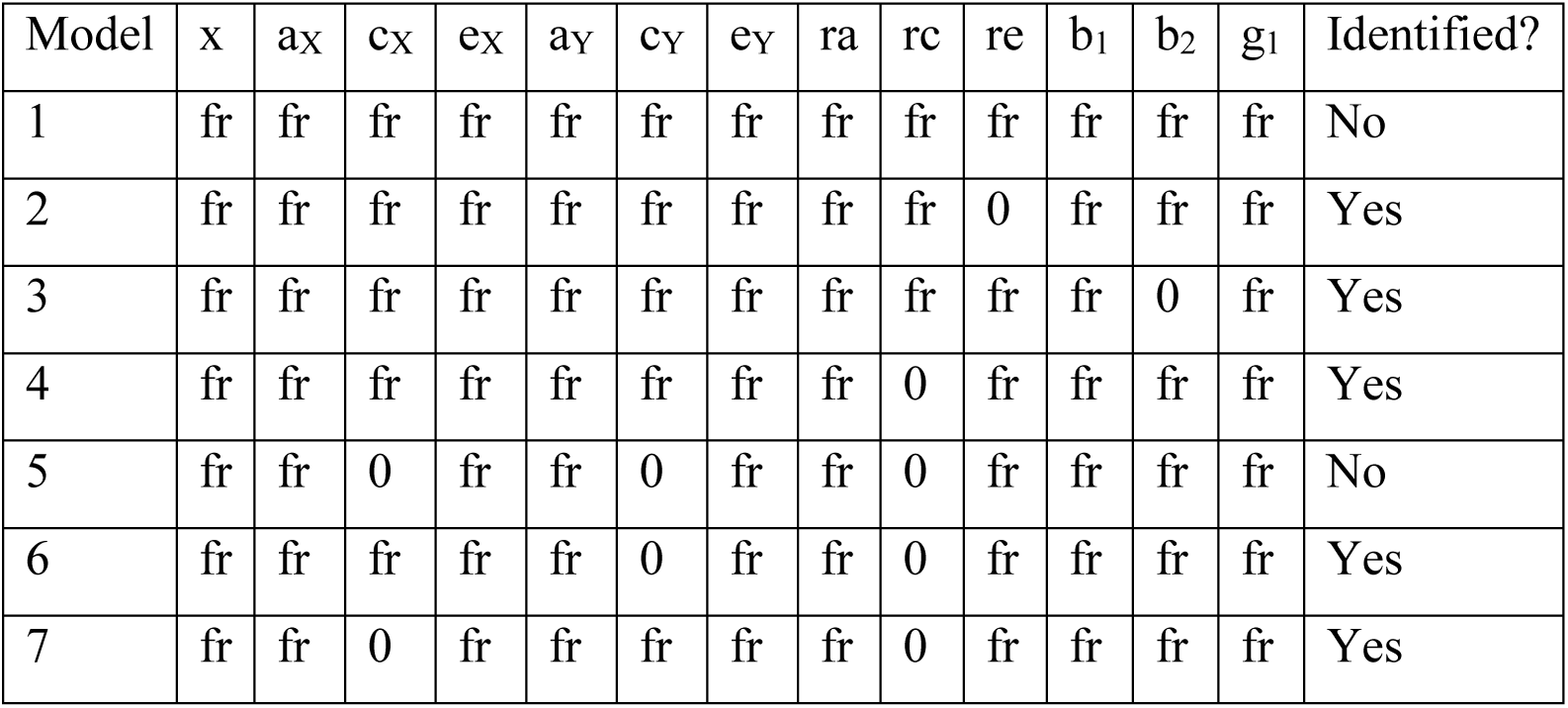
Parameter constraints that render identified the Figure 3. Note ‘fr’ indicates that the parameter is estimated, ‘0’ – that the parameter is constrained to equal 0.

| Model | x | a<sub>x</sub> | c<sub>x</sub> | e<sub>x</sub> | a<sub>y</sub> | c<sub>y</sub> | e<sub>y</sub> | r<sub>a</sub> | r<sub>c</sub> | r<sub>e</sub> | b<sub>1</sub> | b<sub>2</sub> | g<sub>1</sub> | Identified? |
| --- | --- | --- | --- | --- | --- | --- | --- | --- | --- | --- | --- | --- | --- | --- |
| 1 | fr | fr | fr | fr | fr | fr | fr | fr | fr | fr | fr | fr | fr | No |
| 2 | fr | fr | fr | fr | fr | fr | fr | fr | fr | 0 | fr | fr | fr | Yes |
| 3 | fr | fr | fr | fr | fr | fr | fr | fr | fr | fr | fr | 0 | fr | Yes |
| 4 | fr | fr | fr | fr | fr | fr | fr | fr | 0 | fr | fr | fr | fr | Yes |
| 5 | fr | fr | 0 | fr | fr | 0 | fr | fr | 0 | fr | fr | fr | fr | No |
| 6 | fr | fr | fr | fr | fr | 0 | fr | fr | 0 | fr | fr | fr | fr | Yes |
| 7 | fr | fr | 0 | fr | fr | fr | fr | fr | 0 | fr | fr | fr | fr | Yes |

The MR-DoC model is based on the following regression model:

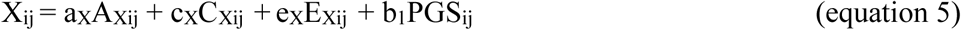

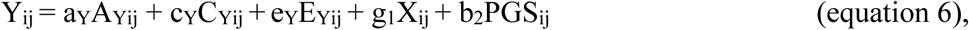

where i stands for twin pairs, and j for twin (j=1,2). The vector of parameters is **θ** = [ra, rc, re, a_x_, c_x_, e_x_, a_y_, c_y_, e_y_, g_1_, b_1_, b_2_, x], where x is the standard deviation of the PGS. Here, and in Figure 3, the parameter of interest is g1, as it concerns the causal effect of exposure X on outcome Y. Note that this model now includes the parameter b_2_, i.e., the pleiotropic effects of PGS on Y, which are usually assumed to be absent. Where we refer to a pleiotropic instrument below, we mean that the parameter b_2_ is not zero. Using ML estimation, we can test hypotheses concerning the parameters by means of a likelihood ratio or Wald test.

### Model identification

Model identification concerns the question whether the observed data provide sufficient information to yield unique estimates of the unknown parameters collected in the vector **θ** ^51^. In the present case, the observed information is summarized in the MZ and DZ (6x6) covariance matrices. We assume that the means are equal in the MZ and DZ and are equal in the twin 1 and twin 2 members (this is obviously testable). As the parameterization of the means has no bearing on the identification of the covariance structure model, we do not consider them in addressing identification.

Local identification is evaluated at a given set of parameters, say **θ_a_,** and implies there are no points in the vicinity of point **θ_a_** in the parameter space leading to the same expected covariance matrices Σ_MZ_(**θ_a_**) and Σ_DZ_(**θ_a_**) ^51,52^. We evaluated local identification using symbolic algebra in Maple ^53^. Derks et al. ^54^ previously used this method in the context of twin modeling. Using Maple, we checked whether the rank of the Jacobian matrix is full column rank. The Jacobian matrix contains the first order derivatives of the elements in the matrices Σ_MZ_(**θ_a_**) and Σ_DZ_(**θ_a_**) with respect to the free parameters. If the Jacobian is not full column rank, we require additional parameter constraints (on the elements in the parameter vector **θ_a_**). Having established local identification in this manner, we proceeded to address the question of resolution by considering the statistical power to estimate the parameters of interest.

### Power calculations

We used exact data simulation ^55^ to create data that fit a given identified model exactly (i.e., the observed covariance matrices equaled the expected covariance matrices exactly). We then dropped parameters of interest and assessed the power in the standard way using the non-central χ^2^ distribution, with noncentrality parameter (NCP) λ. We adopted an alpha of .05 throughout. Data were simulated in R using the MASS library function mvrnorm() with the empirical=TRUE option ^56^. The MR-DoC model was fitted to the population covariance matrices in OpenMx ^57^. We used the R-package AER ^58^ to conduct 2SLS estimation in the standard MR using the sample to twin 1 data, i.e., effectively a sample of unrelated individuals. We used the mRnd power calculator to calculate the power of the 2SLS procedure ^59^. We chose effect sizes by considering the decomposition of the variance of the outcome Y, as illustrated in Figure 4. That is, given that the outcome Y is standardized 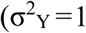), we considered the components making up the explained variance, i.e., 1-σ^2^ξ_Y_, as a function of the regression parameters b_1_, b_2_, g_1_, the variances of the residuals ξ_X_ and ξ_Y_ (parameters 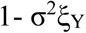 and 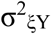), and the covariance between the residuals ξ_X_ and ξ_Y_ (parameter σ_ξXξY_).

We varied (a) the strength of the instrumental variable, defined as the percentage of variance explained in the exposure X by instrument (PGS); (b) the variance of ξ_X_ (residual X), i.e., parameter σ^2^_ξX_, representing the percentage of variance in the exposure, not explained by the instrumental variable); (c) the variance in ξ_Y_ (residual Y), i.e., parameter σ^2^ξ_Y_, representing the percent of variance in Y not explained by the MR model; and (d) the covariance between ξ_X_ and ξ_Y_ (parameter σ_ξXξY_). Using the path tracing rules, we can distinguish five components of variance (C1 to C5, Fig.5) that involve the parameters of interest g_1_ (the causal effect) and b_2_ (the direct effect of the instrument on the outcome). In all scenarios, the predictors explained 10% of the phenotypic variance of the outcome Y. The parameter values used in simulations are included in Supplementary Tables S1 and S2. To provide an indication on the potential gains in power conferred by our approach relative to a standard MR analysis of data obtained in unrelated individuals, we report the number of unrelated persons required to attain equivalent power as MR-DoC based on 2000 twin pairs.

**Figure 5:**
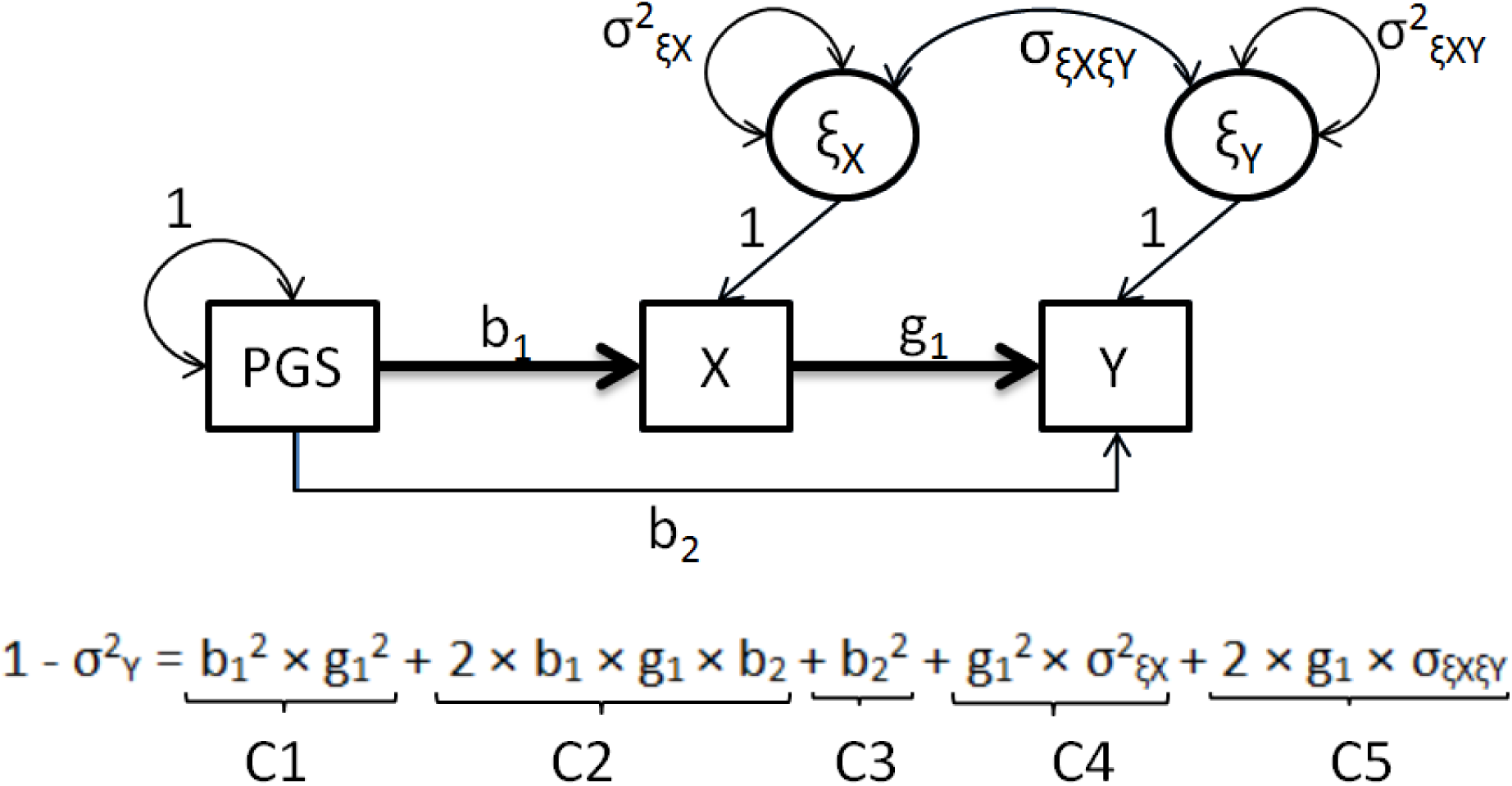
Effect size calculation for the power analyses. Abbreviations: PGS – polygenic score; X – exposure variable; Y – outcome variable; ξ_X_ - residual X; σ^2^_ξX_ - variance in the exposure, not explained by the instrumental variable PGS; ξ_Y_ - residual Y; σ^2^ξ_Y_ - variance in Y not explained by the MR model; σ_ξXξY_ - the covariance between ξ_X_ and ξ_Y_ ; b_1_ – regression coefficient in the regression of the exposure X on the instrument; b_2_ – regression coefficient in the regression of the outcome on the PGS; g_1_ – the causal effect of X on Y.

## RESULTS

### Parameter recovery in standard MR and MR-DoC with valid or invalid (i.e., pleiotropic) instrumental variables (given re = 0)

Table 2 displays results obtained using non-pleiotropic (i.e., Figure 3, b_2_=0), or pleiotropic (i.e., b_2_ ≠ 0, see Fig.4) instrumental variables in the causal effect estimation. Given b_2_=0, results indicate that all estimation methods recover the true parameter value (scenario S1, Table 2). As is to be expected, b_2_≠0 leads to biased estimates of the causal effect when employing standard MR methods (e.g., 2SLS or ratio of coefficients). MR-DoC recovers correctly the true parameter values (scenario S2, Table 2). Finally, we checked parameter recovery when the instrument has pleiotropic effects, but there is no causal effect of the exposure X on outcome Y (i.e., b_2_≠0 and g_1_=0; scenario S3, Table 2). Results showed that standard MR detects a causal effect (when in truth there is none), while MR-DoC does not. We remind the reader that we have set re = 0 to obtain these results.

**Table 2:**
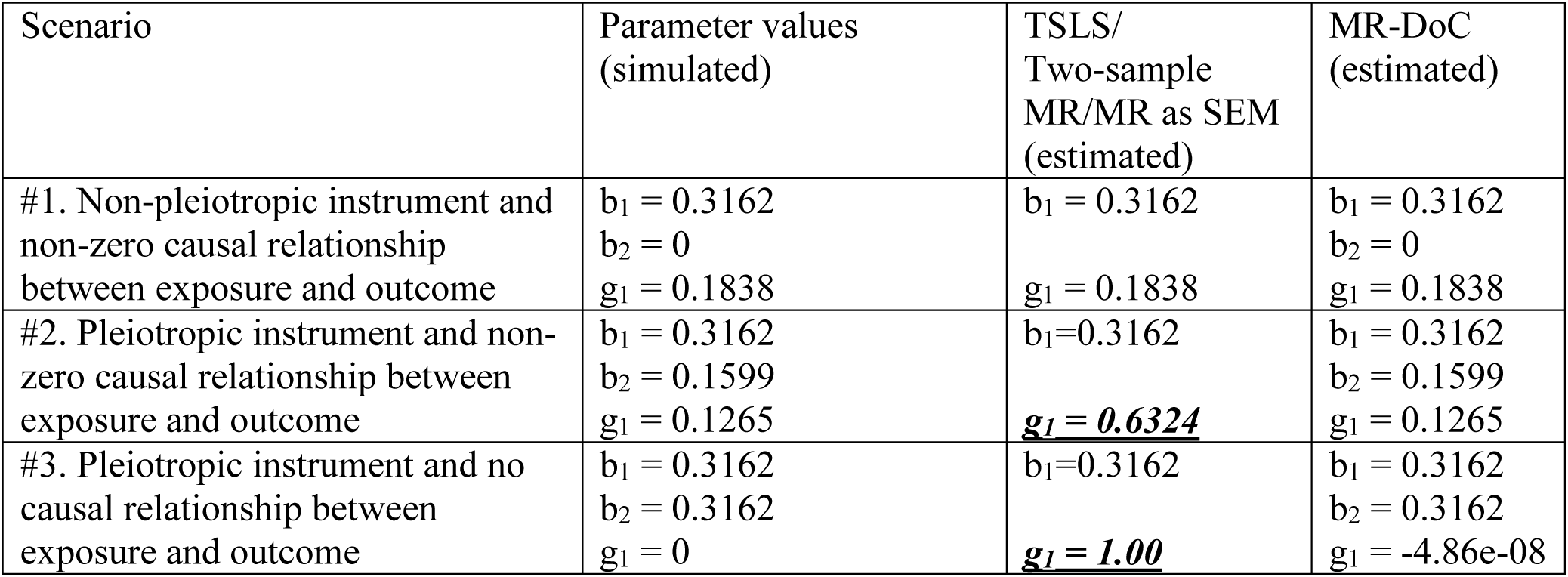
Results of the comparison of the standard MR in analysis of unrelated individuals and MR-DoC twin model. Additive genetic (A), shared environmental (C) and unique environmental (E) effects contributed to the variance of both the exposure 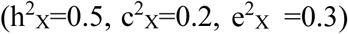 and the outcome 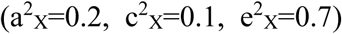 variable, and genetic and shared environmental factors contributed to the correlations between the traits (r_ξXξY_=0.2). Incorrectly estimated parameter values are in italics.

### Model identification

We considered identification in seven models given in Table 1. Model 1, in which all parameters are estimated freely, is not identified. However, constraining to zero any of the parameters re (Model 2), b_2_ (Model 3) or rc (Model 4) renders the model identified (i.e., Models 2, 3, 4). Conversely, all parameters are identified if the two traits differ with respect to their ACE model. This is the case if, e.g., the exposure is an AE trait and the outcome is an ACE trait (implying the parameters c_x_ and rc are zero, Model 7), or the exposure is an ACE trait and the outcome (conditional on the exposure) is an AE trait (implying the parameters c_y_ and rc to zero, Model 6). Note that in the latter case Y (unconditional) is characterized by shared environmental effects on Y (transmitted from C_X_, through the g_1_ path) ^41^. Furthermore, we found that MR-DoC is not identified when the traits’ variances are limited to two sources (e.g., X and Y are both AE traits, Model 5). Fixing the parameter re to 0 is a constraint commonly employed in the fixed economic within MZ twin intra-pair differences model ^44^ and in the discordant twin design. The following results are based on the model with this identifying constraint in place, i.e., re = 0.

### Power

Figure 6 (and Supplementary Tables S3-S5) displays the results pertaining to the power to detect the causal effect in standard MR and in MR-DoC.

**Figure 6:**
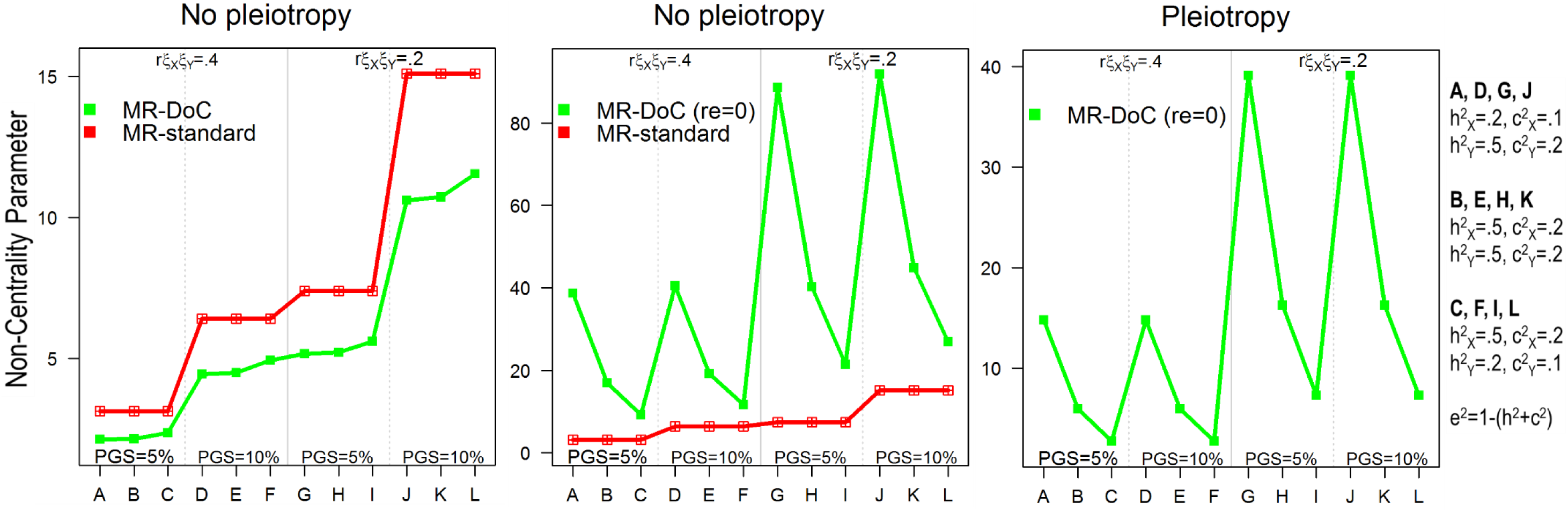
Results given a *non-pleiotropic* instrument (parameter b_2_=0; left and middle panels), and given a *pleiotropic* instrument (parameter b_2_ ≠ 0; right panel). Fitting the model with the parameter g_1_ freely estimated, and with the parameter g_1_ constrained to equal zero provided the Non-Centrality Parameter. The standard MR is based on two-stage least squares in a sample of 4000 unrelateds. The MR-DoC twin model used maximum likelihood and a sample of 2000 twin pairs. Abbreviations: r_ξXξY_ - residual exposure-outcome correlation; PGS – polygenic score.

With a valid instrumental variable (no pleiotropy) and all parameters freely estimated (including the parameter re, Table S3), the main factors that influence MR-DoC’s power are: (a) instrument’s strength; (b) the genetic covariance structure of X (exposure) and Y (outcome); and (c) the magnitude of the residual X-Y correlation. As expected, and consistent with the standard MR literature, increasing instrument’s strength increases power. For instance, with an ACE trait as the exposure variable 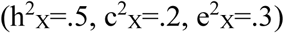, an outcome variable having roughly the same mode of inheritance 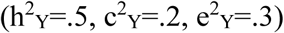, and a residual correlation of r_ξXξY_=.2, the power of the MR-DoC equals .627 and .905 when the instrument explains 5% and 10% of the variance in the exposure, respectively. However, contrary to the standard DoC literature, having traits with similar genetic covariance structure has no bearing on MR-DoC’s power to detect the causal effect. This is the case for instance, if both the outcome and exposure are ACE traits. Power is the highest when the outcome variable has a lower heritability than the exposure variable. For example, power increases from .622 in Scenario S1G (with a 50% heritable outcome and a 20% heritable exposure) to .658 in Scenario S1I (with a 20% heritable outcome variable and a 50% heritable exposure; Table S3). Finally, increasing X-Y residual correlation reduces MR-DoC’s power. For instance, with an outcome and an exposure having roughly the same mode of inheritance, and an instrument explaining 5% of the variance in the exposure, MR-DoC’s power drops from .627 (Scenario S1H, Table S3) to .312 (Scenario S1B) when the residual correlation increases from r_ξXξY_=0.2 to r_ξXξY_=0.4.

Given equal N, standard MR (4000 unrelateds) has larger power than MR-DoC (4000 twins) only when b_2_=0 and re ≠0 (Fig.6, left). Yet, assuming data collected in twins are readily available, greater statistical power is available in the MR-DoC model (than from reducing the twin pairs to singletons and resorting to standard MR, Table S4).

We also calculated the required N of unrelated individuals to achieve the same power as 2000 twin pairs (Table 3). We found the yield of MR-DoC substantial if the parameter re was fixed to zero (as simulated, Fig.6, middle). For example, with a sample of 2000 twin pairs, MR-DoC yields a NCP λ of 38.68 given an instrument explaining 10% of the variance in the exposure, a residual correlation of r_ξXξY_=0.2, an exposure variable with low heritability 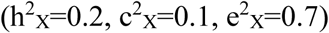, and a moderately heritable outcome 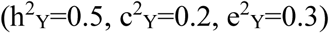. Standard MR needs about 56 737 unrelated individuals to achieve equivalent power (Table 3).

**Table 3:** The number of unrelated individuals (N) required by MR-standard to achieve the same NCP as MR-DoC in 2000 twin pairs. The number of unrelateds was estimated based on the NCP obtained by fitting the standard MR as a structural equation model, with estimation of the causal effect based on maximum likelihood (similar to MR-DoC).

| Scenario (S1) | N required by MR standard to achieve the same NCP as MR-DoC with **re=0** |
| --- | --- |
| A | 56 737 |
| B | 24 945 |
| C | 13 472 |
| D | 28 515 |
| E | 13 480 |
| F | 8206 |
| G | 53 774 |
| H | 24 440 |
| I | 13 013 |
| J | 13 558 |
| K | 13 239 |
| L | 7954 |

Given b_2_≠0 and re=0, MR-DoC’s power increases with: (a) decreasing X-Y residual correlation, and (b) decreasing heritability of the exposure (Fig.6 right; see for details, Tables S2 and S5). Regarding the former, power is always larger when the association between the exposure and the outcome is largely causal in nature (i.e., when the residual correlation drops from r_ξXξY_ = 0.4 to r_ξXξY_ = 0.2). Regarding the latter, power is always greater in scenarios where the exposure has low heritability. For instance, the NCP λ increases from 7.29 to 39.1 when the heritability of the exposure decreases from 50% (S2I) to 20% (S2J, Table S5). The same pattern of results was observed when b_2_=0 and re=0 (see Fig.6 middle, and Table S4). Note that the instrument’s strength has no longer a bearing on power when b2≠0, i.e., when it affects the outcome both directly (parameter b_2_) and indirectly, via the exposure (parameter b_1_). For instance, 2000 twin pairs yield an NCP λ of 5.98 given two traits having identical variance components 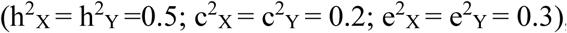, and a large residual correlation (r_ξXξY_ = 0.4), regardless of whether the instrument explains 5% or 10% of the variance in the exposure (Scenarios S2B and S2E, Table S5). As mentioned above, contrary to the standard DoC literature ^43^, MR-DoC does not require the exposure and the outcome variable to have radically different covariance structures to ensure sufficient power to test unidirectional causal hypotheses.

## DISCUSSION

Our aim was to integrate MR and the classical twin model to render testable the MR’s strong assumption that the instrumental variable has no direct effect on the outcome, conditional on the exposure and confounders. We showed that, with standard MR methods, violations of this assumption readily lead to biased causal effect estimates, or may even yield spurious false positives. MR-DoC correctly detects causal effects and provides accurate parameter estimates even if the instrument is pleiotropic. With traits that have the same covariance structure (e.g., when both the outcome and exposure are ACE traits), the weaker assumption (also used in the discordant twin design) that the unique environmental component influencing the exposure, influences the outcome only via its effect on the exposure, but not directly (i.e., re=0), is needed to identify model parameters. Note, however, that this assumption is not required with traits that have different covariance structures (e.g., with an ACE and an AE trait).

Aside from providing the means to relax the ‘no pleiotropy’ assumption, the MR-DoC twin model confers several other advantages in understanding causal relationships between exposures and outcomes. First, MR-DoC provides full statistical description of the observed exposure-outcome relationship, allowing one to disentangle the causal effect of the exposure on the outcome, from potential pleiotropic effects of the instrumental variable, as well as from the contribution of other genetic and environmental factors to the variances and the covariance of the two traits. Second, the twin data provide sufficient information to estimate the direct path from the instrument to the outcome (i.e., the parameter b_2_ in Fig.3). Importantly, this path captures not only pleiotropic effects, but also possible effects of other variants affecting the outcome which are in linkage disequilibrium with the instrument. Third, our approach opens up the possibility of using strong genetic instrumental variables in the form of polygenic scores. This is generally desirable in the standard MR design, as the strength of the valid instrumental variable has a bearing on the precision of the causal estimate (i.e., with weak instruments the estimate tends to approach the observed OLS exposure-outcome association ^60^), and on the distribution of the causal effect estimate (the weaker the instrument, the more skewed the distribution; see Fig.7.1 in ^13^). As a consequence, significance tests and the construction of the confidence intervals, which rely on asymptotic normality, are no longer accurate. Consequently, tests may suffer inflated type I error rates ^61^. In addition, strong instruments are desirable from the perspective of power, as our results showed (consistent with the literature ^61^).

Interestingly, the strength of the instrument (i.e., defined in terms of percentage of explained variance in the exposure), has no bearing on the power when the instrument has pleiotropic effects, given that these effects are correctly modeled (in MR-DoC). The reason for this is that the exposure no longer features as a full mediator variable in the presence of a direct path between the instrument and the outcome. Correspondingly, the misfit due to dropping the causal parameter from the analysis is attenuated by the presence of the parameter b_2_ in the model. That is, fixing the parameter g_1_ to zero will largely bias the parameter b_2_, but will not affect the between twin covariance matrix (as is the case in the standard MR design where the sole path from the instrument to the outcome is via the exposure). Stated otherwise, the bias in b_2_ will be greatest, leaving ra (Ax,Ay) and rc (Cx,Cy) relatively unaffected, regardless of how strong the instrument is. In this circumstance, the power to detect the causal effect will primarily depend on the magnitude of the residual correlation between the two traits and, to a lesser extent, on their modes of inheritance.

MR-DoC is tailored for the readily available datasets collected worldwide on more than 1.5 million twins at the Twin and Family Registries (see ^62^ for details on these rich resources of genotypic and phenotypic data). We showed that twin data correctly detect causation and estimate effect sizes even in the presence of pleiotropy. Although primarily developed to address the ‘no pleiotropy’ assumption, our results demonstrate that MR-DoC has greater statistical power than standard MR analysis in singletons if the parameter re is zero (as fixed to zero in the model). With re fixed to zero, dropping the causal parameter g_1_ from the model greatly impacts all paths connecting the exposure and the outcome, both within and across twins (parameters ra and rc), creating a large discrepancy between the observed and the expected covariance matrix. This misfit is evident throughout the covariance matrix. Consequently, in some specific circumstances, testing causal hypotheses requires tens of thousands of unrelated individuals for one to achieve the same power as that conferred by several hundreds of twin pairs. It should be noted that unlike MR methods ^37,38^ or asymmetry tests ^34^ based on summary statistics, the approach proposed here only requires GWAS results for the exposure variable. With MR-DoC and these rich phenotype resources - ranging from personality, diet and lifestyle to disease and psychiatric traits ^62^ - collected at Twin and Family registries, the availability of genetic instruments robustly associated with the exposure remains the main limiting factor in addressing causal questions in non-experimental settings.

As presented here, MR-DoC is limited to twin data, but note that twin registries often have available information on additional family members ^62-66^. Conversely, not all cohort studies necessarily include related individuals. To expand the model, we aim to accommodate additional family members (siblings and parents), and distantly related individuals by using Genetic Relationship Matrices (GRMs) based on average allelic correlations (where the alleles are defined at the measured single nucleotide polymorphisms). We anticipate that these extensions will further increase statistical power and robustness to assumption violation ^67,68^. Second, throughout the paper we assumed that the mating is random, there is no genotype-environment correlation, and no genotype by environment interaction. Indeed, assumption violation may also arise because the mating is assortative ^13^, or because there are other plausible paths from the instrument to the outcome (except direct paths, or indirect, via the exposure), for example, via confounders affecting both traits, i.e., implying genotype by environment interaction or genotype by environment correlation. However, we note that these effects may be captured by the MR-DoC twin model with appropriate experimental designs ^67,69^. On a cautionary note, although valid strong instruments are desirable in MR from the perspective of power, making up the polygenic score based on variants of unknown function should limit the testable hypotheses to whether the model is consistent or not with a direct causal effect from the exposure to the outcome (as pointed out in ^21^). While we considered the use of MR-DoC with polygenic scores, our conclusions also hold in scenarios where a genetic variant with known function is used as the instrument, which would improve the biological interpretation of the causal effect.

In conclusion, by integrating Mendelian Randomization with the Direction of Causation twin model, we developed a model that allows one to test and relax the strong ‘no pleiotropy’ assumption employed by standard MR. This approach therefore allows one to employ strong instrumental variables in the form of polygenic scores, guarding against the weak instrument bias and increasing the power to detect causal effects of exposures on potential outcomes. We anticipate that MR-DoC will enhance and extend MR’s range of applications, and increase the value of the large cohorts collected at twin registries as they correctly detect causation and estimate effect sizes even in the presence of pleiotropy.

## Supplementary Tables

**Table S1:** Scenarios and parameter values used to generate data for assessing the power of the MR–DoC twin model, given a *non-pleiotropic* instrumental variable (parameter b_2_≠0). We varied the strength of the instrument, (i.e., the Polygenic Score explained either 0.05 or 0.1 proportion of the variance in the exposure), the proportion of variance explained in the exposure (X) and the outcome (Y) variable by additive genetic 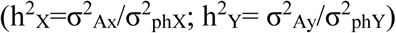, shared environmental 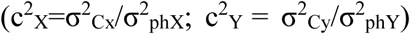 and unique environmental 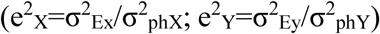 factors, and the contribution of the instrumental variable (PGS; parameter b_1_), the causal effect (g_1_) and of the residual correlation between the outcome and the exposure (rξXξY) to the 10% explained variance in the outcome. The effect size is defined as the percentage of explained variance in the outcome given the chosen parameter values b_1_, b_2_, g_1_, σ_ξXξY_ (covariance of ξX and ξY), σ^2^_ξX_ (residual variance in X), and σ^2^_ξY_ (residual variance in Y).

| Scenario (S1) | Parameter values | Effect size: percentage of variance explained in the outcome (%) |
| --- | --- | --- |
| A | PGS = 0.05<br>$h^2_X = 0.2$, $c^2_X = 0.1$, $e^2_X = 0.7$<br>$h^2_Y = 0.5$, $c^2_Y = 0.2$, $e^2_Y = 0.3$<br>$r_{\xi X \xi Y} = 0.4$ | $b_1^2 \times g_1^2 = 0.068$<br>$g_1^2 \times \sigma^2_{\xi X} = 1.295$<br>$2 \times g_1 \times \sigma_{\xi X \xi Y} = 8.636$ |
| B | PGS = 0.05<br>$h^2_X = 0.5$, $c^2_X = 0.2$, $e^2_X = 0.3$<br>$h^2_Y = 0.5$, $c^2_Y = 0.2$, $e^2_Y = 0.3$<br>$r_{\xi X \xi Y} = 0.4$ | $b_1^2 \times g_1^2 = 0.068$<br>$g_1^2 \times \sigma^2_{\xi X} = 1.295$<br>$2 \times g_1 \times \sigma_{\xi X \xi Y} = 8.636$ |
| C | PGS = 0.05<br>$h^2_X = 0.5$, $c^2_X = 0.2$, $e^2_X = 0.3$<br>$h^2_Y = 0.2$, $c^2_Y = 0.1$, $e^2_Y = 0.7$<br>$r_{\xi X \xi Y} = 0.4$ | $b_1^2 \times g_1^2 = 0.068$<br>$g_1^2 \times \sigma^2_{\xi X} = 1.295$<br>$2 \times g_1 \times \sigma_{\xi X \xi Y} = 8.636$ |
| D | PGS = 0.1<br>$h^2_X = 0.2$, $c^2_X = 0.1$, $e^2_X = 0.7$<br>$h^2_Y = 0.5$, $c^2_Y = 0.2$, $e^2_Y = 0.3$<br>$r_{\xi X \xi Y} = 0.4$ | $b_1^2 \times g_1^2 = 0.142$<br>$g_1^2 \times \sigma^2_{\xi X} = 1.278$<br>$2 \times g_1 \times \sigma_{\xi X \xi Y} = 8.579$ |
| E | PGS = 0.1<br>$h^2_X = 0.5$, $c^2_X = 0.2$, $e^2_X = 0.3$<br>$h^2_Y = 0.5$, $c^2_Y = 0.2$, $e^2_Y = 0.3$<br>$r_{\xi X \xi Y} = 0.4$ | $b_1^2 \times g_1^2 = 0.142$<br>$g_1^2 \times \sigma^2_{\xi X} = 1.278$<br>$2 \times g_1 \times \sigma_{\xi X \xi Y} = 8.579$ |
| F | PGS = 0.1<br>$h^2_X = 0.5$, $c^2_X = 0.2$, $e^2_X = 0.3$<br>$h^2_Y = 0.2$, $c^2_Y = 0.1$, $e^2_Y = 0.7$<br>$r_{\xi X \xi Y} = 0.4$ | $b_1^2 \times g_1^2 = 0.142$<br>$g_1^2 \times \sigma^2_{\xi X} = 1.278$<br>$2 \times g_1 \times \sigma_{\xi X \xi Y} = 8.579$ |
| G | PGS = 0.05<br>$h^2_X = 0.2$, $c^2_X = 0.1$, $e^2_X = 0.7$ | $b_1^2 \times g_1^2 = 0.164$<br>$g_1^2 \times \sigma^2_{\xi X} = 3.126$ |
| | $h^2_Y = 0.5, c^2_Y = 0.2, e^2_Y = 0.3$<br>$r_{\xi X \xi Y} = 0.2$ | $2 \times g_1 \times \sigma_{\xi X \xi Y} = 6.70$ |
| H | PGS = 0.05<br>$h^2_X = 0.5, c^2_X = 0.2, e^2_X = 0.3$<br>$h^2_Y = 0.5, c^2_Y = 0.2, e^2_Y = 0.3$<br>$r_{\xi X \xi Y} = 0.2$ | $b_1^2 \times g_1^2 = 0.164$<br>$g_1^2 \times \sigma_{\xi X}^2 = 3.126$<br>$2 \times g_1 \times \sigma_{\xi X \xi Y} = 6.709$ |
| I | PGS = 0.05<br>$h^2_X = 0.5, c^2_X = 0.2, e^2_X = 0.3$<br>$h^2_Y = 0.2, c^2_Y = 0.1, e^2_Y = 0.7$<br>$r_{\xi X \xi Y} = 0.2$ | $b_1^2 \times g_1^2 = 0.164$<br>$g_1^2 \times \sigma_{\xi X}^2 = 3.126$<br>$2 \times g_1 \times \sigma_{\xi X \xi Y} = 6.709$ |
| J | PGS = 0.1<br>$h^2_X = 0.2, c^2_X = 0.1, e^2_X = 0.7$<br>$h^2_Y = 0.5, c^2_Y = 0.2, e^2_Y = 0.3$<br>$r_{\xi X \xi Y} = 0.2$ | $b_1^2 \times g_1^2 = 0.338$<br>$g_1^2 \times \sigma_{\xi X}^2 = 3.042$<br>$2 \times g_1 \times \sigma_{\xi X \xi Y} = 6.619$ |
| K | PGS = 0.1<br>$h^2_X = 0.5, c^2_X = 0.2, e^2_X = 0.3$<br>$h^2_Y = 0.5, c^2_Y = 0.2, e^2_Y = 0.3$<br>$r_{\xi X \xi Y} = 0.2$ | $b_1^2 \times g_1^2 = 0.338$<br>$g_1^2 \times \sigma_{\xi X}^2 = 3.042$<br>$2 \times g_1 \times \sigma_{\xi X \xi Y} = 6.619$ |
| L | PGS = 0.1<br>$h^2_X = 0.5, c^2_X = 0.2, e^2_X = 0.3$<br>$h^2_Y = 0.2, c^2_Y = 0.1, e^2_Y = 0.7$<br>$r_{\xi X \xi Y} = 0.2$ | $b_1^2 \times g_1^2 = 0.338$<br>$g_1^2 \times \sigma_{\xi X}^2 = 3.042$<br>$2 \times g_1 \times \sigma_{\xi X \xi Y} = 6.619$ |

**Table S2:**
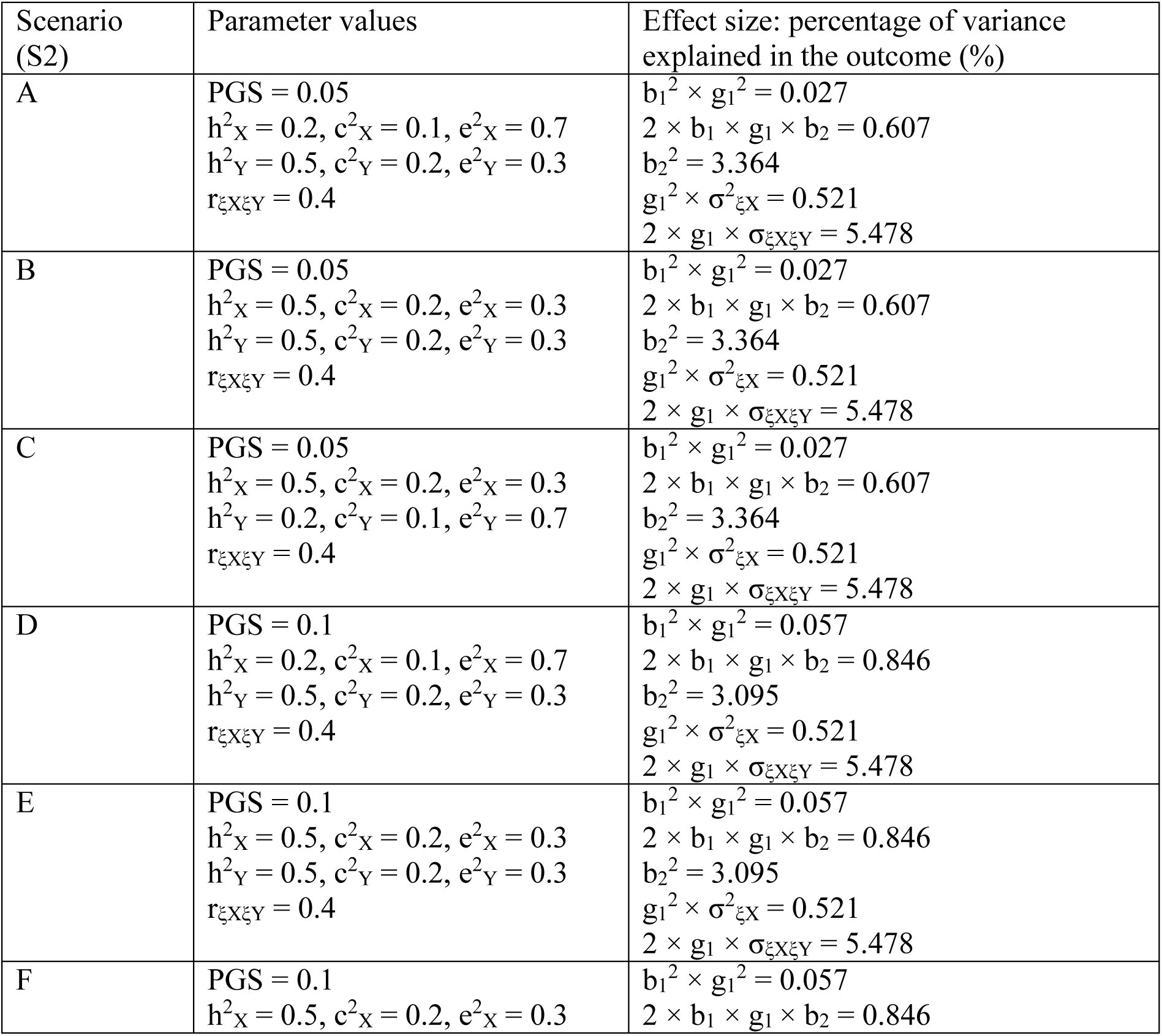
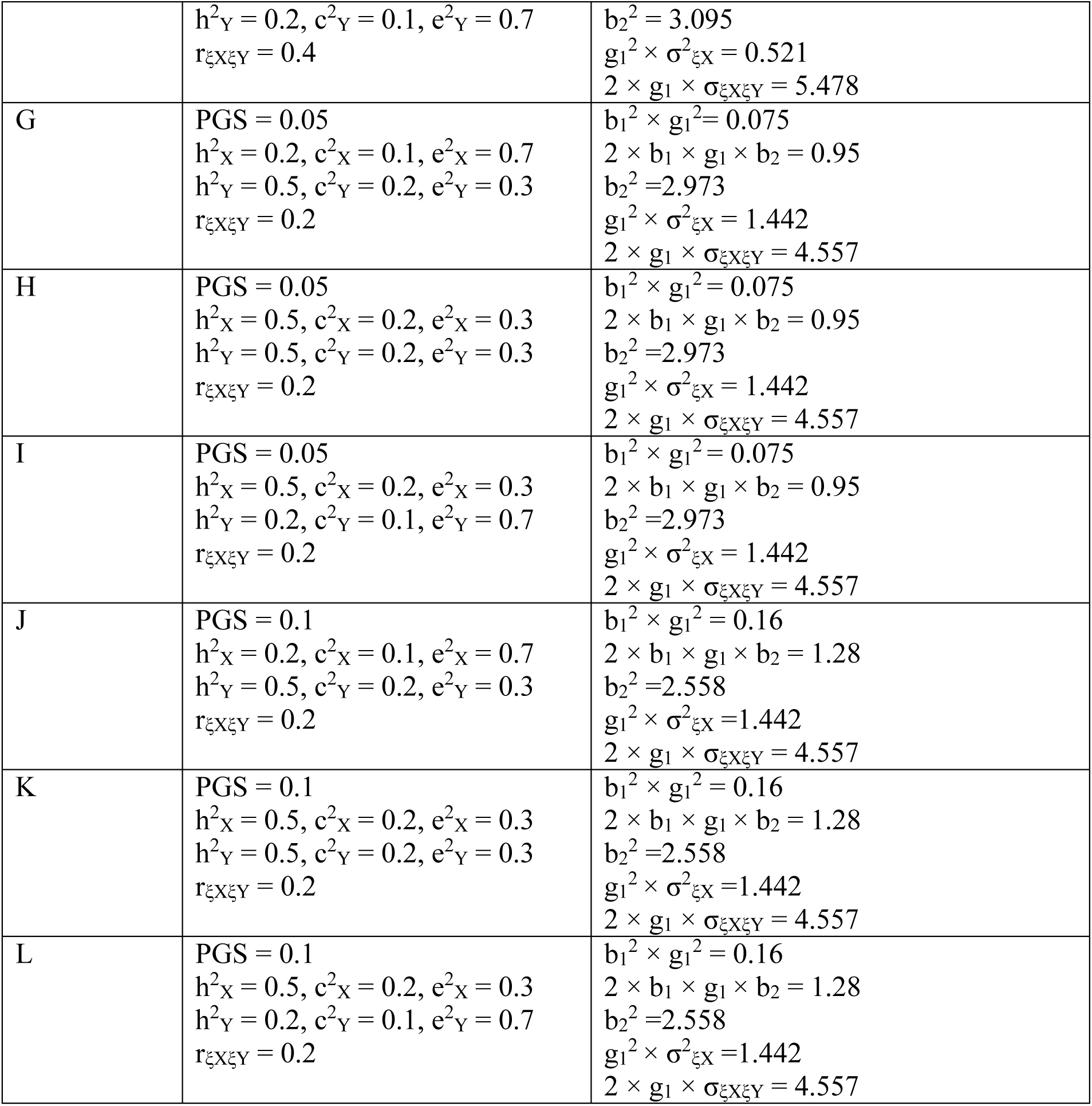
Scenarios and parameter values used to generate data for assessing the power of the MR–DoC twin model, given a *pleiotropic* instrumental variable and no unique environmental correlation (parameters b_2_≠0 and re=0). We varied the strength of the instrument, (i.e., the Polygenic Score explained either 0.05 or 0.1 proportion of the variance in the exposure), the proportion of variance explained in the exposure (X) and the outcome (Y) variable by additive genetic 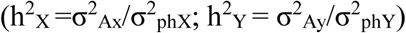, shared environmental 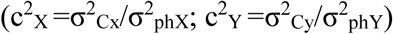 and unique environmental 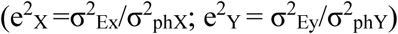 factors, and the contribution of the instrumental variable (PGS; parameter b_1_), the causal effect (g_1_) and of the residual correlation between the outcome and the exposure (rξXξY) to the 10% explained variance in the outcome. The effect size is defined as the percentage of explained variance in the outcome given the chosen parameter values b_1_, b_2_, g_1_, σ_ξXξY_ (covariance of ξX and ξY), σ^2^_ξX_ (residual variance in X), and σ^2^_ξY_ (residual variance in Y).

| Scenario (S2) | Parameter values | Effect size: percentage of variance explained in the outcome (%) |
| --- | --- | --- |
| A | PGS = 0.05<br>$h^2_X = 0.2$, $c^2_X = 0.1$, $e^2_X = 0.7$<br>$h^2_Y = 0.5$, $c^2_Y = 0.2$, $e^2_Y = 0.3$<br>$r_{\xi X \xi Y} = 0.4$ | $b_1^2 \times g_1^2 = 0.027$<br>$2 \times b_1 \times g_1 \times b_2 = 0.607$<br>$b_2^2 = 3.364$<br>$g_1^2 \times \sigma^2_{\xi X} = 0.521$<br>$2 \times g_1 \times \sigma_{\xi X \xi Y} = 5.478$ |
| B | PGS = 0.05<br>$h^2_X = 0.5$, $c^2_X = 0.2$, $e^2_X = 0.3$<br>$h^2_Y = 0.5$, $c^2_Y = 0.2$, $e^2_Y = 0.3$<br>$r_{\xi X \xi Y} = 0.4$ | $b_1^2 \times g_1^2 = 0.027$<br>$2 \times b_1 \times g_1 \times b_2 = 0.607$<br>$b_2^2 = 3.364$<br>$g_1^2 \times \sigma^2_{\xi X} = 0.521$<br>$2 \times g_1 \times \sigma_{\xi X \xi Y} = 5.478$ |
| C | PGS = 0.05<br>$h^2_X = 0.5$, $c^2_X = 0.2$, $e^2_X = 0.3$<br>$h^2_Y = 0.2$, $c^2_Y = 0.1$, $e^2_Y = 0.7$<br>$r_{\xi X \xi Y} = 0.4$ | $b_1^2 \times g_1^2 = 0.027$<br>$2 \times b_1 \times g_1 \times b_2 = 0.607$<br>$b_2^2 = 3.364$<br>$g_1^2 \times \sigma^2_{\xi X} = 0.521$<br>$2 \times g_1 \times \sigma_{\xi X \xi Y} = 5.478$ |
| D | PGS = 0.1<br>$h^2_X = 0.2$, $c^2_X = 0.1$, $e^2_X = 0.7$<br>$h^2_Y = 0.5$, $c^2_Y = 0.2$, $e^2_Y = 0.3$<br>$r_{\xi X \xi Y} = 0.4$ | $b_1^2 \times g_1^2 = 0.057$<br>$2 \times b_1 \times g_1 \times b_2 = 0.846$<br>$b_2^2 = 3.095$<br>$g_1^2 \times \sigma^2_{\xi X} = 0.521$<br>$2 \times g_1 \times \sigma_{\xi X \xi Y} = 5.478$ |
| E | PGS = 0.1<br>$h^2_X = 0.5$, $c^2_X = 0.2$, $e^2_X = 0.3$<br>$h^2_Y = 0.5$, $c^2_Y = 0.2$, $e^2_Y = 0.3$<br>$r_{\xi X \xi Y} = 0.4$ | $b_1^2 \times g_1^2 = 0.057$<br>$2 \times b_1 \times g_1 \times b_2 = 0.846$<br>$b_2^2 = 3.095$<br>$g_1^2 \times \sigma^2_{\xi X} = 0.521$<br>$2 \times g_1 \times \sigma_{\xi X \xi Y} = 5.478$ |
| F | PGS = 0.1<br>$h^2_X = 0.5$, $c^2_X = 0.2$, $e^2_X = 0.3$ | $b_1^2 \times g_1^2 = 0.057$<br>$2 \times b_1 \times g_1 \times b_2 = 0.846$ |
| | $h^2_Y = 0.2, c^2_Y = 0.1, e^2_Y = 0.7$<br>$r_{\xi X \xi Y} = 0.4$ | $b_2^2 = 3.095$<br>$g_1^2 \times \sigma^2_{\xi X} = 0.521$<br>$2 \times g_1 \times \sigma_{\xi X \xi Y} = 5.478$ |
| G | $PGS = 0.05$<br>$h^2_X = 0.2, c^2_X = 0.1, e^2_X = 0.7$<br>$h^2_Y = 0.5, c^2_Y = 0.2, e^2_Y = 0.3$<br>$r_{\xi X \xi Y} = 0.2$ | $b_1^2 \times g_1^2 = 0.075$<br>$2 \times b_1 \times g_1 \times b_2 = 0.95$<br>$b_2^2 = 2.973$<br>$g_1^2 \times \sigma^2_{\xi X} = 1.442$<br>$2 \times g_1 \times \sigma_{\xi X \xi Y} = 4.557$ |
| H | $PGS = 0.05$<br>$h^2_X = 0.5, c^2_X = 0.2, e^2_X = 0.3$<br>$h^2_Y = 0.5, c^2_Y = 0.2, e^2_Y = 0.3$<br>$r_{\xi X \xi Y} = 0.2$ | $b_1^2 \times g_1^2 = 0.075$<br>$2 \times b_1 \times g_1 \times b_2 = 0.95$<br>$b_2^2 = 2.973$<br>$g_1^2 \times \sigma^2_{\xi X} = 1.442$<br>$2 \times g_1 \times \sigma_{\xi X \xi Y} = 4.557$ |
| I | $PGS = 0.05$<br>$h^2_X = 0.5, c^2_X = 0.2, e^2_X = 0.3$<br>$h^2_Y = 0.2, c^2_Y = 0.1, e^2_Y = 0.7$<br>$r_{\xi X \xi Y} = 0.2$ | $b_1^2 \times g_1^2 = 0.075$<br>$2 \times b_1 \times g_1 \times b_2 = 0.95$<br>$b_2^2 = 2.973$<br>$g_1^2 \times \sigma^2_{\xi X} = 1.442$<br>$2 \times g_1 \times \sigma_{\xi X \xi Y} = 4.557$ |
| J | $PGS = 0.1$<br>$h^2_X = 0.2, c^2_X = 0.1, e^2_X = 0.7$<br>$h^2_Y = 0.5, c^2_Y = 0.2, e^2_Y = 0.3$<br>$r_{\xi X \xi Y} = 0.2$ | $b_1^2 \times g_1^2 = 0.16$<br>$2 \times b_1 \times g_1 \times b_2 = 1.28$<br>$b_2^2 = 2.558$<br>$g_1^2 \times \sigma^2_{\xi X} = 1.442$<br>$2 \times g_1 \times \sigma_{\xi X \xi Y} = 4.557$ |
| K | $PGS = 0.1$<br>$h^2_X = 0.5, c^2_X = 0.2, e^2_X = 0.3$<br>$h^2_Y = 0.5, c^2_Y = 0.2, e^2_Y = 0.3$<br>$r_{\xi X \xi Y} = 0.2$ | $b_1^2 \times g_1^2 = 0.16$<br>$2 \times b_1 \times g_1 \times b_2 = 1.28$<br>$b_2^2 = 2.558$<br>$g_1^2 \times \sigma^2_{\xi X} = 1.442$<br>$2 \times g_1 \times \sigma_{\xi X \xi Y} = 4.557$ |
| L | $PGS = 0.1$<br>$h^2_X = 0.5, c^2_X = 0.2, e^2_X = 0.3$<br>$h^2_Y = 0.2, c^2_Y = 0.1, e^2_Y = 0.7$<br>$r_{\xi X \xi Y} = 0.2$ | $b_1^2 \times g_1^2 = 0.16$<br>$2 \times b_1 \times g_1 \times b_2 = 1.28$<br>$b_2^2 = 2.558$<br>$g_1^2 \times \sigma^2_{\xi X} = 1.442$<br>$2 \times g_1 \times \sigma_{\xi X \xi Y} = 4.557$ |

**Table S3:** Simulation results given a *non-pleiotropic* instrumental variable (parameter b_2_ = 0). We report the power to detect the causal effect g_1_ (given alpha=0.05), and the NCP obtained based on the standard MR with the causal effect estimated using two-stage least squares (N=2000 unrelateds), and based on the MR-DoC twin model (N=2000 twin pairs). In fitting the MR-DoC model we estimate all model parameters (with the parameter re, simulated re=0, freely estimated).

| Scenario (S1) | MR-DoC's power (NCP) in 2000 twin pairs | Standard MR's power (NCP) in 2000 unrelated individuals (twin1) | Standard MR's power (NCP) in 4000 unrelated individuals |
| --- | --- | --- | --- |
| A | .309 (2.13) | .25 (1.61) | .42 (3.13) |
| B | .312 (2.15) | .25 (1.61) | .42 (3.13) |
| C | .336 (2.36) | .25 (1.61) | .42 (3.13) |
| D | .559 (4.44) | .44 (3.25) | .72 (6.41) |
| E | .564 (4.50) | .44 (3.25) | .72 (6.41) |
| F | .602 (4.93) | .44 (3.25) | .72 (6.41) |
| G | .622 (5.16) | .49 (3.73) | .78 (7.39) |
| H | .627 (5.21) | .49 (3.73) | .78 (7.39) |
| I | .658 (5.61) | .49 (3.73) | .78 (7.39) |
| J | .902(10.615) | .79 (7.59) | .97 (15.1) |
| K | .905 (10.72) | .79 (7.59) | .97 (15.1) |
| L | .924 (11.54) | .79 (7.59) | .97 (15.1) |

**Table S4:** Simulation results given a *non-pleiotropic* instrumental variable (b_2_=0). We report the power to detect the causal effect g_1_ (given alpha of 0.05) and the NCP obtained based on the standard MR with the causal effect estimated using two-stage least squares (N=2000 unrelateds), and based on the MR-DoC twin model (N=2000 twin pairs). In fitting the MR-DoC model we constrained re to equal 0 (as simulated).

| Scenario (S1) | Power (NCP) based on 2000 twin-pairs | Standard MR's power (NCP) based on 2000 unrelateds (twin 1) | Standard MR's power (NCP) in 4000 unrelateds |
| --- | --- | --- | --- |
| A | >.99 (38.68) | .25 (1.61) | .42 (3.13) |
| B | .984 (17.00) | .25 (1.61) | .42 (3.13) |
| C | .857 (9.18) | .25 (1.61) | .42 (3.13) |
| D | >.99 (40.52) | .44 (3.25) | .72 (6.41) |
| E | .992 (19.15) | .44 (3.25) | .72 (6.41) |
| F | .927 (11.66) | .44 (3.25) | .72 (6.41) |
| G | >.99 (88.54) | .45 (3.73) | .78 (7.39) |
| H | >.99 (40.24) | .45 (3.73) | .78 (7.39) |
| I | >.99 (21.42) | .45 (3.73) | .78 (7.39) |
| J | >.99 (91.83) | .79 (7.59) | .97 (15.1) |
| K | >.99 (44.83) | .79 (7.59) | .97 (15.1) |
| L | >.99 (26.93) | .79 (7.59) | .97 (15.1) |

**Table S5:** Simulation results given a *pleiotropic* instrumental variable (parameter b_2_ ≠ 0). We report the power to detect the causal effect g_1_ (given alpha=0.05), and the NCP based on the MR-DoC twin model (N=2000 twin pairs). In fitting the MR-DoC model, to render the model identified, we assumed that parameter re equals 0 (as simulated).

| Scenario (S2) | Power (NCP) |
| --- | --- |
| A | .97 (14.77) |
| B | .686 (5.98) |
| C | .379 (2.73) |
| D | .970 (14.77) |
| E | .686 (5.98) |
| F | .379 (2.73) |
| G | >.99 (39.1) |
| H | .981 (16.28) |
| I | .770 (7.29) |
| J | >.99 (39.1) |
| K | .981 (16.28) |
| L | .77 (7.29) |

